# Ciliation of muscle stem cells is critical to maintain regenerative capacity and is lost during aging

**DOI:** 10.1101/2020.03.20.000943

**Authors:** A.R. Palla, K.I. Hilgendorf, A.V. Yang, J.P. Kerr, A.C. Hinken, J. Demeter, P. Kraft, N.A. Mooney, N. Yucel, P.K. Jackson, H.M. Blau

## Abstract

During aging, the regenerative capacity of muscle stem cells (MuSCs) decreases, diminishing the ability of muscle to repair following injury. We performed a small molecule library screen and discovered that the proliferation and expansion of aged MuSCs is regulated by signal transduction pathways organized by the primary cilium, a cellular protrusion that serves as a sensitive sensory organelle. Abolishing MuSC cilia *in vivo* severely impaired injury-induced muscle regeneration. In aged muscle, a cell intrinsic defect in MuSC ciliation leading to impaired Hedgehog signaling was associated with the decrease in regenerative capacity. This deficit could be overcome by exogenous activation of Hedgehog signaling which promoted MuSC expansion, both *in vitro* and *in vivo*. Delivery of the small molecule Smoothened agonist (SAG) to muscles of aged mice restored regenerative capacity leading to increased strength post-injury. These findings provide fresh insights into the signaling dysfunction in aging and identify the ciliary Hedgehog signaling pathway as a potential therapeutic target to counter the loss of muscle regenerative capacity which accompanies aging.

## Introduction

Adult muscle stem cells (MuSCs) are crucial for skeletal muscle regeneration throughout life. Sarcopenia, the age-dependent loss of skeletal muscle mass and strength, is a major public-health problem that affects an estimated 15% of individuals 65 years or older^1, 2^. With aging, MuSCs undergo intrinsic changes that reduce their number and function, which decreases the capacity of muscle to respond to exercise and injury and efficiently repair damage to muscle fibers. Such MuSC intrinsic changes include atypical activation of p38-MAPK or Jak2-Stat3 signaling^3, 4, 5, 6, 7^. Additionally, extrinsic changes that alter the niche, or MuSC microenvironment, impact MuSC regenerative function with aging^4^. Finally, maintenance of the quiescent cell state is essential for muscle stem cell function^8^, and the increase in cytokines in aged muscles, including FGF2, WNT3A and TGF-β, alters the signaling state of aged MuSCs and promotes their premature exit from quiescence^4, 8^. Despite this knowledge, our understanding of the basis for MuSC dysfunction with aging remains incomplete.

Many adult stem cells possess primary cilia, an antenna-like structure that protrudes from the cell surface to mediate sensory and morphogenetic signaling^9^. Primary cilia transduce signaling from a growing list of G-protein coupled receptors (GPCRs), proteins with seven transmembrane helices capable of transducing extracellular stimuli into intracellular signals. GPCRs are of particular interest as they constitute a potent class of drug targets^10^. A recent report showed that MuSCs possess a primary cilium that is linked to their cell cycle state^11^, but the role of primary cilia in MuSC function *in vivo* has yet to be established. Here, we show for the first time that genetic ablation of primary cilia in adult MuSCs dramatically decreases their self-renewal and regenerative capacity *in vivo*. Additionally, we identify loss of ciliation as an intrinsic defect of MuSCs with aging, and that augmenting the regenerative capacity of aged MuSCs through activation of the ciliary Hedgehog (Hh) signaling pathway is a previously unrecognized therapeutic strategy for the treatment of sarcopenia.

## Results

### Drug screen to identify novel regulators of aged MuSC proliferation yields cilia-related hits

Aged MuSCs are known to have reduced proliferative capacity which correlates with reduced regeneration of muscle post-injury. To identify intrinsic pathways that could improve the regenerative capacity of muscle in the aged, we designed a screen to test the effect of a curated list of approximately 200 small molecules targeting GPCRs on proliferation of aged MuSCs. We isolated and cultured α7^+^CD34^+^Lin^−^ MuSCs, as previously described^12^, from aged mice (24-26 mo.) **(Fig. 1A)**. Aged MuSCs were exposed to compounds 24 hr post-plating and their proliferative capacity assessed 7 days later using VisionBlue, a Resazurin-based dye that becomes fluorescent upon reduction by metabolically active cells. We confirmed that the VisionBlue signal correlated with cell number in control experiments (**Fig. S1A**), and included previously identified compounds known to promote MuSC proliferation, the prostaglandin PGE2^13^ and the p38-MAPK inhibitor SB202190^5^, as positive controls. (**Fig. S1B**). Cell density was calculated (**Fig. S1C**) and the data normalized to vehicle-treated MuSCs in the same plate. Using this screening strategy, we identified 11 compounds that had an effect on the proliferation of aged MuSCs **(Fig. 1B-D)**. Notably, almost half of these compounds targeted GPCRs localized to the primary cilium^10^ **(Fig. 1C, D)**. Among them was an agonist of the Smoothened (SMO) receptor that robustly induced proliferation of aged MuSCs **(Fig. 1D)**. To confirm our finding, we treated aged MuSCs with known agonists of the SMO receptor, including three small molecules (SAG, purmorphamine and GSA-10)^14^, the glucocorticoid fluticasone^15^, and one of its activating ligands, the protein Sonic Hedgehog (Shh)^16^. We found that these SMO agonists also promoted aged MuSC proliferation **(Fig. 1E)**. Since Sonic Hedgehog is known to signal through the primary cilium^17^, we hypothesized that ciliary signaling could play an important role in activating the regenerative capacity of MuSCs.

**Figure 1.**
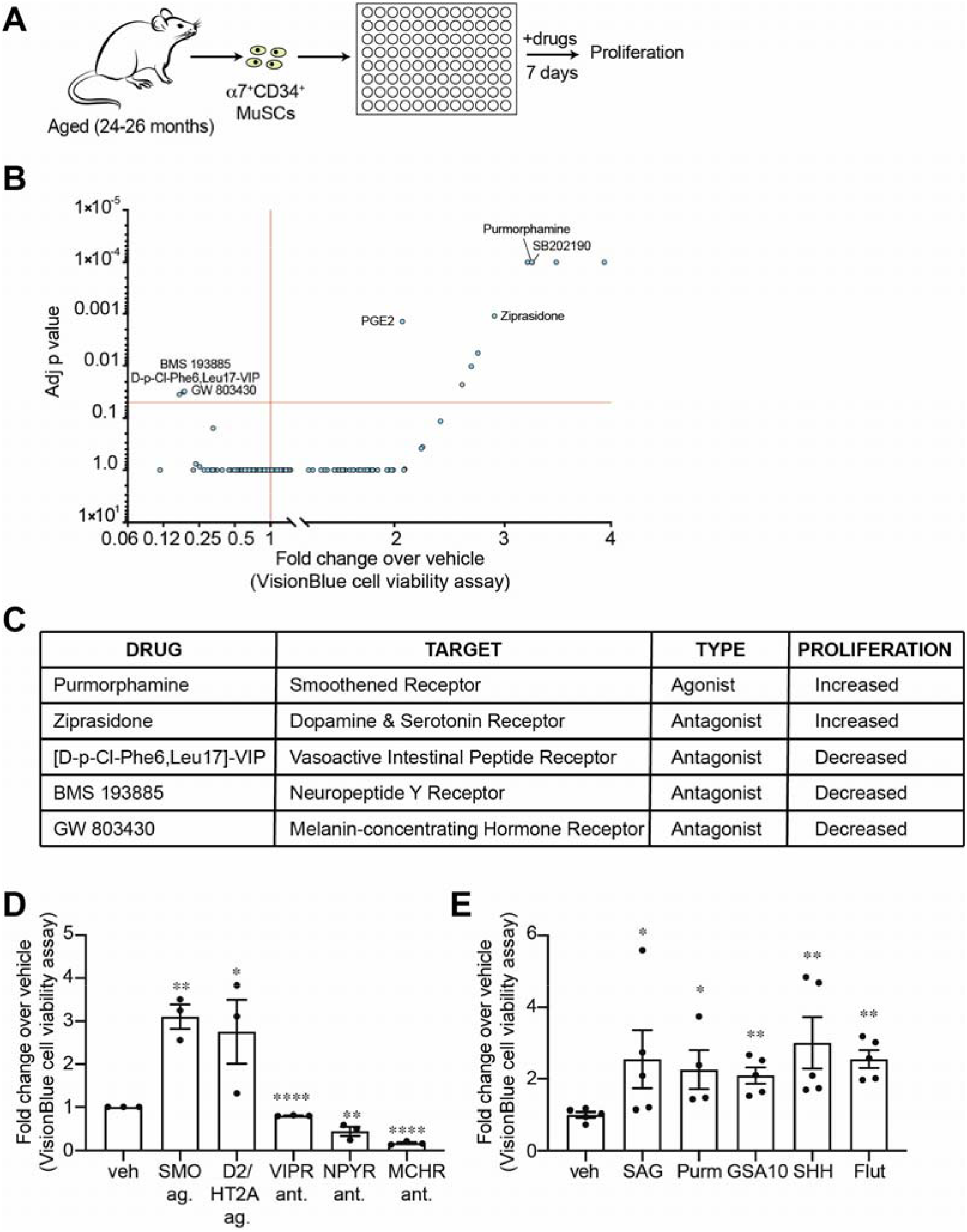
Drug screen in aged MuSCs identifies ciliary GPCRs as regulators of proliferation. **(A)** MuSCs (α7+CD34+) were isolated from aged hindlimb muscles (24-26 months) and treated in culture with a library of compounds (10 μM) targeting G protein-coupled receptors (GPCRs). Proliferation was measured 7 days post-plating (n=5 biological replicates/condition). **(B)** Volcano plot showing p-values and fold change of proliferation of treated normalized to vehicle. **(C)** Significant (p<0.05) ciliary GPCR targets identified to modulate aged MuSC proliferation. **(D)** Proliferation of aged MuSCs treated with ciliary GPCR targeting drugs, shown as fold change normalized to vehicle-treated. SMO, Smoothened Receptor; D2/HT2A, Dopamine&Serotonin Receptor; VIPR, Vasoactive Intestinal Peptide Receptor; NPYR, Neuropeptide Y Receptor; MCHR, Melanin-concentrating Hormone Receptor. **(E)** Proliferation of aged MuSCs treated with SMO agonist, shown as fold change normalized to vehicle-treated (Purm, Purmorphamine; SHH, Sonic Hedgehog; Flut, Fluticasone). *P<0.05, **P<0.01, ***P<0.001 ****P<0.0001. Student’s t-test performed for each condition compared to its vehicle treated control. Means+s.e.m.

### Loss of cilia on MuSCs impairs muscle regeneration and strength recovery

We sought to determine if primary cilia in MuSCs are required for their self-renewal and regenerative capacity in response to injury. Pax7 is the hallmark transcription factor expressed by MuSCs^18^. We therefore generated a Pax7^CreERT2^;IFT88^fl/fl^ mouse model in which the intraflagellar transport protein IFT88, required for ciliary assembly and maintenance^19^, is specifically and conditionally ablated in Pax7-expressing MuSCs (**Fig. 2A**). We confirmed a decrease in the number and length of cilia in tamoxifen-treated Pax7^CreERT2^;IFT88^fl/fl^ (IFT88^−/−^) MuSCs compared to littermate Pax7^CreERT2^;IFT88^+/+^ controls (control) on isolated myofibers by immunofluorescence microscopy (**Fig. 2B,C**). Of note, heterozygous Pax7^CreERT2^;IFT88^+/fl^ (IFT88^+/−^) MuSCs exhibited an intermediate decrease in ciliation (**Fig. 2B)**. To assess the functional consequence of MuSC ciliation loss, we performed notexin-induced injury in the *Gastrocnemius* muscle of young (10-12 weeks) control IFT88^+/−^ and IFT88^−/−^ mice and measured strength recovery. We found that homozygous loss of IFT88 led to a 50% decrease in tetanic force 7 days post-injury and a 20% decrease at 14 days post-injury compared to controls (**Fig. 2D**, **Fig. S2A,B**). Similarly, muscle mass was decreased in IFT88^−/−^ mice at day 14 (**Fig. S2C)**. Impaired regenerative capacity was corroborated by an increase in number of myofibers with reduced cross-sectional area (CSA) at day 14 post-injury quantified by immunofluorescence (**Fig. 2E-G**). The observed intermediate decrease in regenerative capacity of IFT88^+/−^ MuSCs is consistent with the partial loss of ciliation observed in MuSCs from heterozygous mice (Fig 2B). These results demonstrate that the primary cilium of MuSCs plays an important role in notexin-induced muscle regeneration.

**Figure 2.**
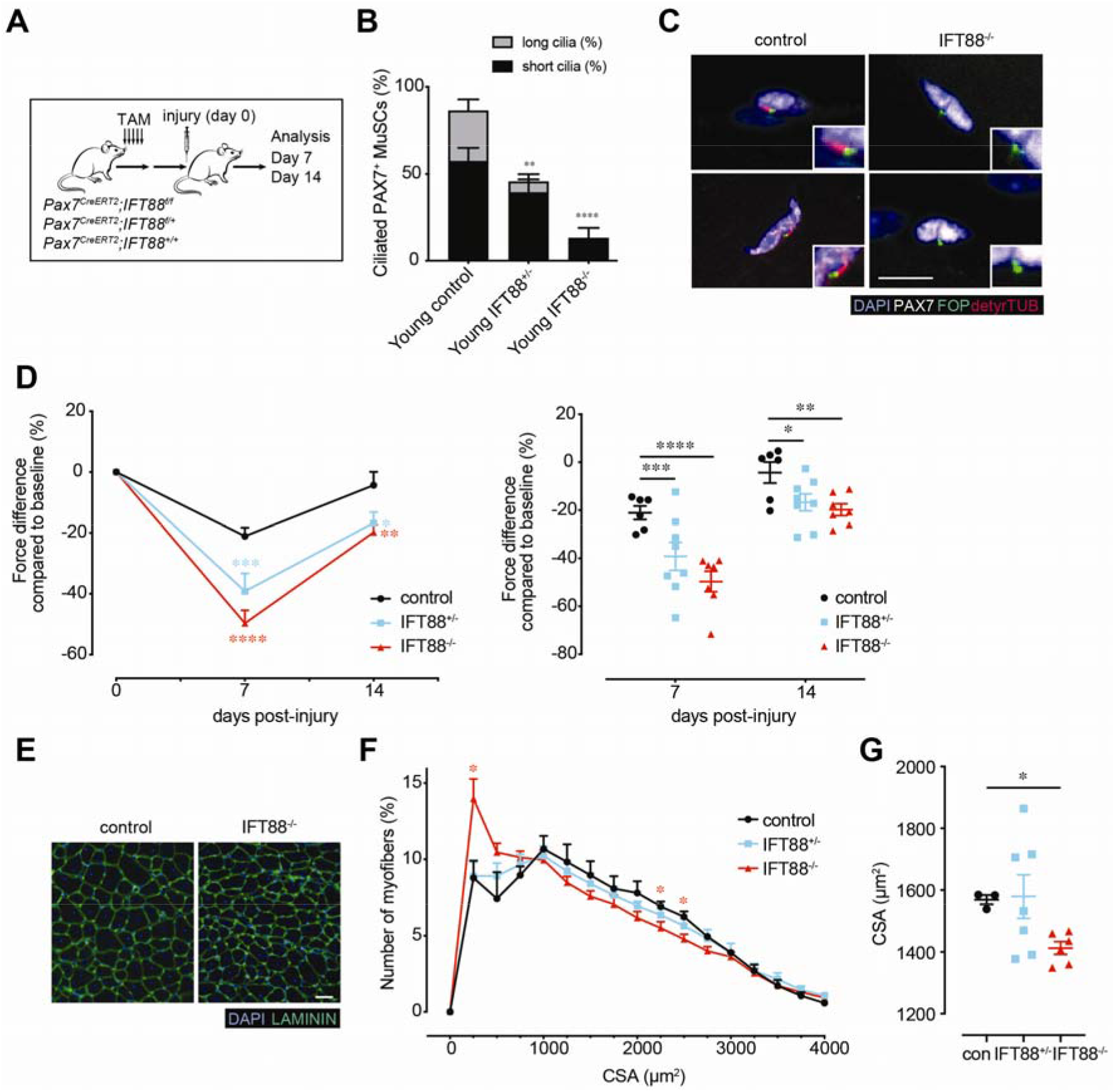
Loss of primary cilia in MuSCs impairs muscle regeneration and strength recovery. **(A)** Pax7-specific Ift88 conditional knockout mice (*Pax7*^*CreERT2*^*;IFT*^−/−^, IFT88^−/−^), heterozygous mice (*Pax7*^*CreERT2*^*;IFT*^+/*f*^, IFT88^+/−^) or control littermates (*Pax7*^*CreERT2*^*;IFT88*^+/+^, control) were treated with tamoxifen (TAM) at 8 weeks of age, injured with notexin and analysis was performed 7 or 14 days post-injury (n=3 control mice; n=4 heterozygous and knockout mice). **(B)** Percent of short (<1 μM) and long (>1 μM) cilia on Pax7+ MuSCs quantified from isolated myofibers of control, IFT88^+/−^ and IFT88^−/−^ mice (n >30 myofibers from each mouse, n=4 mice per genotype). **(C)** Representative confocal images of uninjured/resting EDL myofibers of control and IFT88^−/−^ mice showing cilia immunostaining in Pax7^+^ MuSCs. Scale bars: 10 μm. DAPI, blue; PAX7, white; FOP, green; detyrosinated tubulin, red. **(D)** Plantar flexion tetanic torque of control, IFT88^+/−^ and IFT88^−/−^ mice on day 7 and day 14 post-injury (values normalized to baseline torque). **(E)** Representative Gastrocnemius (GA) cross-section at 14 days post-injury from control and IFT88^−/−^ mice. DAPI, blue; LAMININ, green. Bar=50 μm. **(F)** Myofiber cross-sectional areas (CSA) in control, IFT88^+/−^ and IFT88^−/−^ aged GAs (n=3 for control, n=6 for IFT88^+/−^ and n=6 for IFT88^−/−^). **(G)** Mean CSA. *P<0.05, **P<0.01, ***P<0.001 ****P<0.0001. ANOVA test with Bonferroni correction for multiple comparisons (**B,D,G,H**). Means+s.e.m.

### Primary cilia in MuSCs are required to promote MuSC proliferation and engraftment

To understand if primary cilia are a key intrinsic property of self-renewing MuSCs, we analyzed the regenerative capacity of isolated MuSCs lacking cilia. We analyzed the proliferative potential of MuSCs plated on elastic hydrogels with a stiffness equivalent to that of muscle tissue (12 kPa), which preserves their self-renewal properties in culture^20^. IFT88^−/−^ MuSCs have a reduced proliferative capacity compared to controls, as determined by total number of MuSCs after 7 days (**Fig. 3A**). We further confirmed this reduction in cell division capacity using an EdU incorporation assay (**Fig. 3B**). RNAseq analyses of control and IFT88^−/−^ MuSCs revealed an enrichment of pathways related to calcium signaling (i.e*. Akap5, Cacna1d, Cacna1g, Cacnab, Camk1d, Camkk2, Chrna4, Chrnb2*), Gα signaling (i.e*. Add1, Cnga1, Gnb5, Prkar1a, Rapgef4, Rapgef4, Rgs2*) and cell cycle (i.e*. Atm, Chek2, Babam1, Cdk13, Mcm4*) in IFT88^−/−^ MuSCs (**Fig. 3C**, **S3A, Table S1**) in good agreement with the function of primary cilia in other cell types^21, 22, 23, 24^.

**Figure 3.**
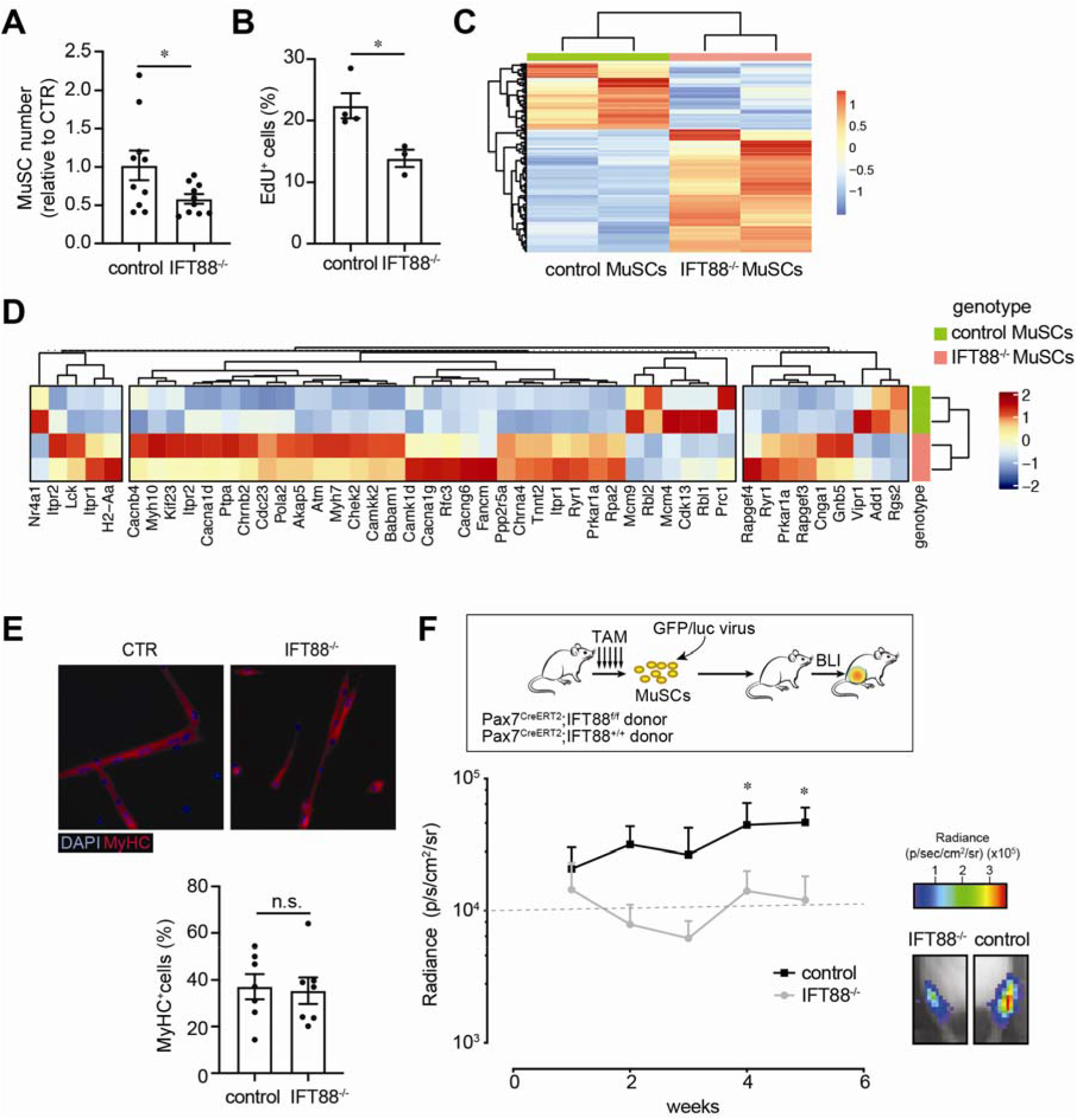
Loss of primary cilia in MuSCs reduces proliferative and self-renewal capacity. **(A)** Number of control and IFT88^−/−^ MuSCs after culture on hydrogel for 7 days, normalized to control. (n=10 per group performed in 3 independent experiments). **(B)** Percentage of EdU^+^ MuSCs (n=4 mice for control, n=3 mice for IFT88^−/−^). **(C)** Heat map of differentially expressed genes (p<0.1) between control or IFT88^−/−^ MuSCs freshly isolated from hindlimb muscles. (**D)** Heat map of genes identified genes in IPA enriched pathways. **(E)** Top: Representative images of cultured control and IFT88^−/−^ MuSCs after 7 days on collagen-coated plates showing MyHC immunostaining. Scale bars: 20 μM. Bottom: Quantification of MyHC positive control and IFT88^−/−^ differentiated MuSCs. **(F)** Engraftment of GFP/luc-labeled control and IFT88^−/−^ MuSCs. Transplant scheme (top). BLI signals post-transplant expressed as average radiance (p◻s^−1^◻cm^−2^◻sr^−1^). (n=5 replicates per condition) (bottom). Representative BLI images for each condition. Bar=5 mm. (bottom-right). Mann-Whitney test (**A, B, E**). ANOVA test for group comparison and significant difference for the endpoint by Fisher’s test (**F).** *P<0.05, **P<0.01. Means+s.e.m.

To elucidate if loss of ciliary signaling and consequent reduction in MuSC proliferation could result from spontaneous differentiation of MuSCs, we plated control and IFT88^−/−^ MuSCs on collagen coated plates. The IFT88^−/−^ MuSCs did not differ from ciliated controls in their spontaneous differentiation and exhibited a comparable capacity to form MyHC positive myotubes (**Fig. 3D**), lending credence to our hypothesis that the role of cilia is in maintaining MuSC proliferation and expansion. To determine if self-renewal capacity was altered *in vivo*, we performed engraftment studies using control and IFT88^−/−^ MuSCs transduced with a GFP/luciferase expression vector, which enables the dynamics of MuSC engraftment and contribution to regeneration to be assessed over time by non-invasive bioluminescence imaging (BLI). NSG mice were irradiated to deplete competing endogenous MuSCs. Equal numbers of GFP/luc expressing Control and IFT88^−/−^ MuSCs were then injected into the *Tibialis anterior* (TA) muscles and engraftment potential measured over time using BLI, as previously described^5, 12, 13^ (**Fig. 3F**). We found that IFT88^−/−^ MuSC engraftment was significantly reduced compared to controls (**Fig. 3F**). These data show that cilia are crucial to the self-renewal and expansion capacity of MuSCs.

### Smoothened receptor agonist (SAG) promotes MuSC expansion in vivo

We tested if activation of the GPCR Smoothened (SMO), which is restricted to cilia^25^, could promote MuSC proliferation. We found that treatment of control, but not IFT88^−/−^ MuSCs with a Smoothened agonist (SAG) led to a robust increase in proliferation (**Fig. 4A**), implicating the Hh signaling pathway in cilia-dependent MuSC expansion. In accordance, inhibition of Hh signaling by cyclopamine^26^ markedly reduced proliferation, as assessed by EdU incorporation (**Fig. 4B)**. To further study if stimulation of the Hedgehog pathway had a beneficial effect on muscle regeneration in vivo, we used a transgenic mouse model we previously developed, Pax7^CreERT2^;Rosa26-LSL-Luc to monitor the dynamics of endogenous MuSC expansion over time using BLI^13^. We found that a single intramuscular injection of SAG significantly increased endogenous MuSC expansion post-injury (**Fig. 4C,D)**. These results demonstrate that ciliary signaling via SMO is important for the proliferative capacity of MuSCs, and that stimulating SMO can lead to increased MuSC engraftment during regeneration.

**Figure 4.**
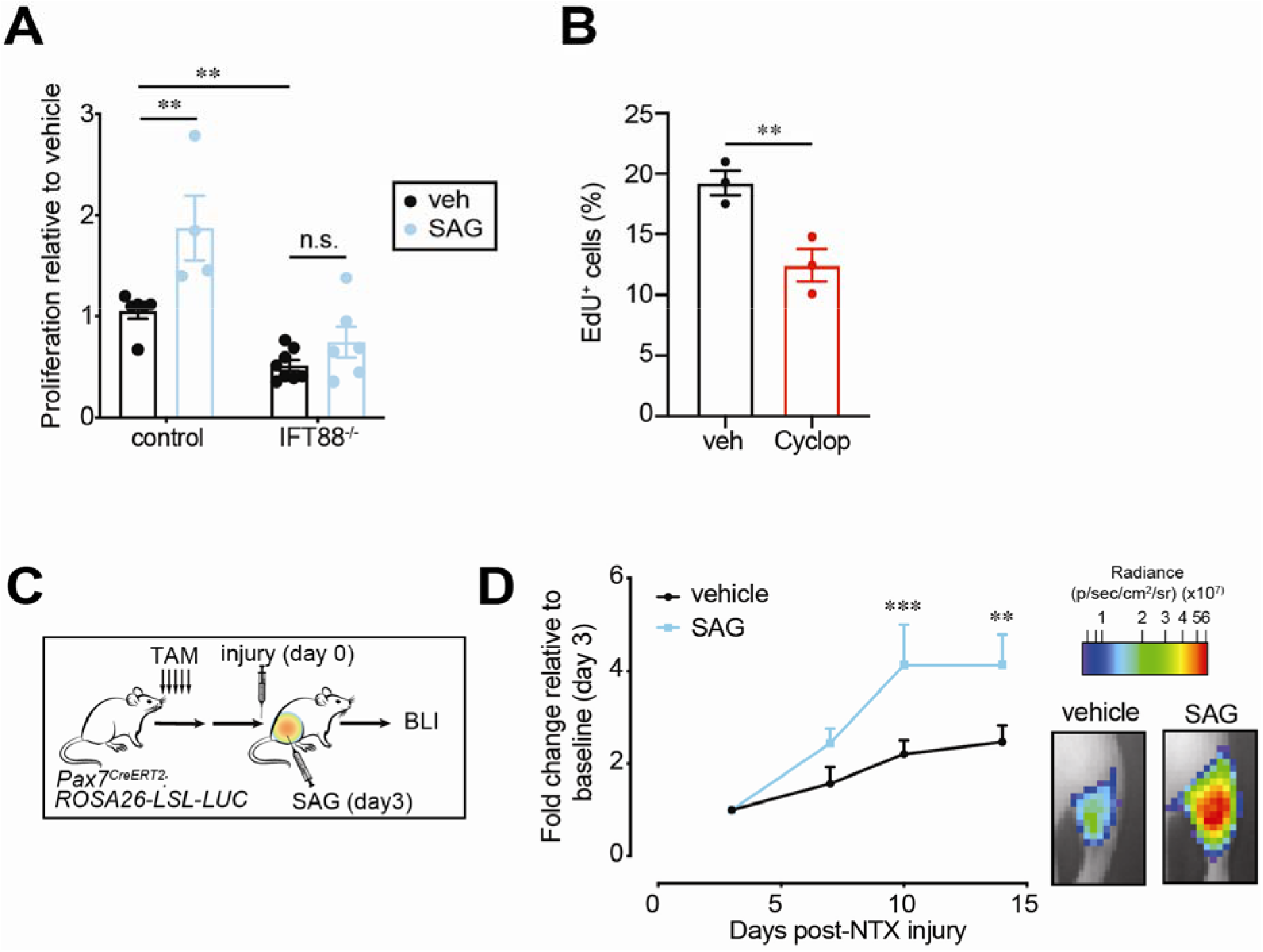
SMO agonist treatment promotes MuSC expansion. **(A)** Proliferation of control or IFT88^−/−^ MuSCs treated with vehicle or SMO agonist SAG, shown as fold change normalized to vehicle-treated. **(B)** Percentage of EdU^+^ vehicle or cyclopamine (SMO inhibitor) treated MuSCs (n=3 mice per condition). **(C,D)** Expansion of endogenous MuSCs in Pax7CreERT2;Rosa26-LSL-Luc mice treated with tamoxifen (TAM) to label resting MuSCs and assayed by BLI post-notexin injury (n = 3 mice per condition). (C) Experimental scheme. (D) Left: BLI (n = 3 mice per condition). Right: Representative BLI image. (Scale bar, 5 mm). *P<0.05, **P<0.01, ***P<0.001 ****P<0.0001. ANOVA test with Bonferroni correction for multiple comparisons (**A**). Mann-Whitney test (**B).** ANOVA test for group comparisons and significant difference for endpoint by Fisher’s test (**D**) Means+s.e.m.

### Aged MuSCs exhibit decreased ciliation and expression of Hh downstream target Gli2

Since ablation of cilia leads to a loss of MuSC self-renewal capacity, we hypothesized that aged MuSCs possess a ciliation defect that contributes to their loss of stemness and regenerative function. We therefore assessed the ciliation status of MuSCs during aging. First, we compared the cilia of young and aged MuSCs associated with isolated myofibers. By immunofluorescence analysis, we identified the cilia of PAX7-positive MuSCs using antibodies directed against detyrosinated tubulin and the centrosomal marker FOP (**Fig. 5A**). Aged MuSCs had a decrease in the number of cilia compared to young (**Fig. 5B**). Upon analysis of transcriptomics data from young and aged MuSCs, we identified several differentially expressed ciliary genes (i.e. *Ift122, Kif3a, Dzip1l, Stil*) (**Fig. 5C**), and in accordance with our data, members of the Hedgehog signaling (Hh) pathway (i.e. *Ptch, Gli2*), known to signal through the primary cilium^17^, were among the top hits (**Fig. 5C**). Moreover, expression of both the SMO receptor and the downstream effector, Gli2, was dysregulated in aged compared to young MuSCs, not only at a basal state, but also post-injury. Unlike in young mice, where these Hh-associated genes exhibit a transient upregulation at day 3 post-injury, aged mouse muscles show a downregulation of SMO and Gli2 following notexin injury (**Fig. 5D**). These data demonstrate that a loss of ciliation in aged MuSCs likely accounts for the observed dysregulation of the Hh signaling pathway.

**Figure 5.**
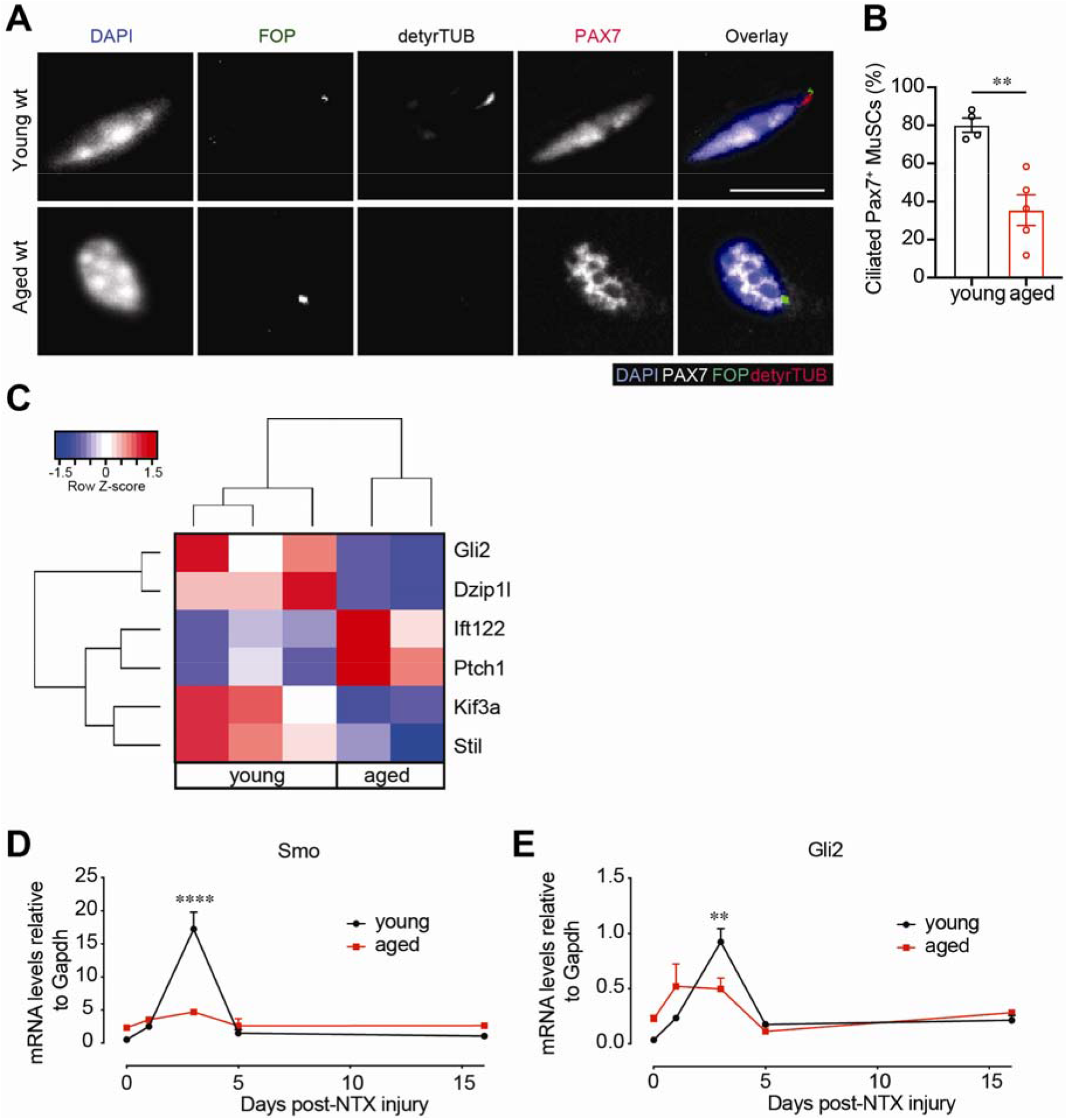
Aged MuSCs present decreased ciliation and Hedgehog signaling. **(A)** Representative confocal images of uninjured/resting EDL myofibers of young (2 months) and aged (25 months) mice (n >30 myofibers from each mouse, n=4 mice for young and 5 mice for aged) showing cilia immunostaining in Pax7^+^ MuSCs. Scale bars: 10 μm. DAPI, blue; PAX7, white; FOP, green; detyrosinated tubulin, red. **(B)** Percent of ciliated Pax7+ MuSCs quantified from isolated myofibers. **(C)** Heat map of differentially expressed HH pathway genes in young and aged MuSCs. **(D,E)** Expression of SMO receptor (**D**) and Gli2 transcription factor (**E**) after TA muscle injury (notexin) (n = 4 mice). *P<0.05, **P<0.01, ***P<0.001 ****P<0.0001. Mann-Whitney test (**B).** ANOVA test for group comparisons and significant difference for each timepoint by Fisher’s test **(D,E).** Means+s.e.m.

### SMO agonist (SAG) promotes young and aged MuSC proliferation and function

To assess if stimulation of Hedgehog signaling in the remaining ~30% ciliated aged MuSCs suffices to improve muscle regeneration in the aged, we treated isolated MuSCs from aged mice with SAG. SAG treatment resulted in a significant increase in proliferation (**Fig. 6A)**, and in the expression of the downstream Hedgehog target Gli2 (**Fig. 6B)**. We performed RNAseq analyses of aged vehicle and SAG-treated MuSCs (**Fig. 6C)**. Using Gene Ontology analyses of the differentially expressed genes between SAG and vehicle treated aged MuSCs, we found an enrichment of ciliary pathways (i.e. *Mkks*), cell cycle (i.e. *Pcid2, Ercc2, Igf2, Kntc1, Stat5b*) and calcium signaling pathways (i.e. *Actc1, Atp2b3, Myl4*) (**Fig. S4, S5, Table S2**). To determine if SAG injection could enhance the function of endogenous aged MuSCs, we injected SAG into the Gastrocnemius (GA) muscles post-injury. Notably, we observed a significant increase in the strength and muscle mass of SAG-treated aged muscles post-injury (**Fig. 6D-F**). Thus, a stimulation of ciliary Hedgehog signaling in aged MuSCs significantly enhances recovery of muscle mass and strength post-injury.

**Figure 6.**
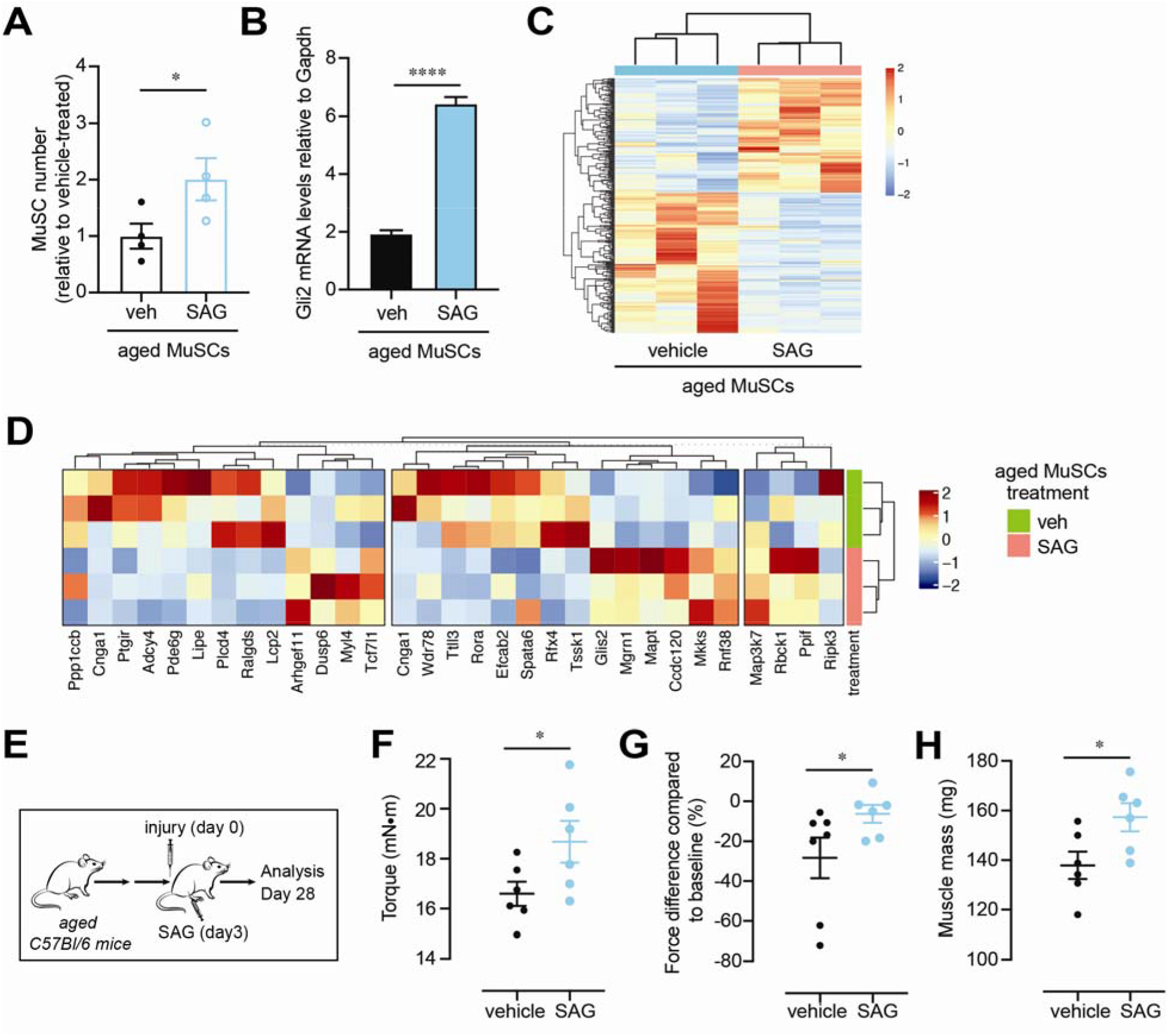
Smo agonist promotes MuSC proliferation and function in aged MuSCs. **(A)** Proliferation of aged MuSCs (>24 mo) treated with vehicle or SAG, shown as fold change normalized to vehicle-treated (n=4 mice per condition). **(B)** Expression levels of SMO downstream target Gli2 in MuSCs 24 hours after vehicle or SAG treatment (n=3 mice per condition). **(C)** Heat map of differentially expressed genes (p<0.1) between vehicle and SAG treated MuSCs treated for 24 hours cultured on hydrogels. (**D)** Heat map of genes identified genes in IPA enriched pathways. **(E-H)** Aged (>24 months) mice were injected with vehicle or SAG post-notexin (NTX) injury and muscle regeneration was analyzed 4 weeks later as force recovery. (**E**) Experimental scheme. (**F**) Plantar flexion tetanic torque (absolute values) (n = 6 mice per condition). (**G**) Plantar flexion tetanic torque relative to baseline (n = 6 mice per condition). (**H**) Gastrocnemius muscle mass (n = 6 mice per condition). *P<0.05, **P<0.01, ***P<0.001 ****P<0.0001. Paired t-test (**A,B).** Mann-Whitney test **(F-H).** Mean + SEM.

## Discussion

Primary cilia are cellular protrusions found on most mammalian cells that serve as sensory organelles that detect diverse signals such as light, growth factors, and morphogens^27^. Perturbation of ciliary proteins leads to a broad range of genetic disorders known as “ciliopathies”, which can affect a numerous cell types and tissues. Hypotonia and muscle flaccidity is one common clinical manifestation, for example in Joubert Syndrome. This is at least in part a consequence of disrupted motor coordination^28^. Here, we uncover a role for primary cilia on muscle stem cells in mediating the Hedgehog signaling pathway to promote muscle regeneration *in vivo* and provide novel evidence of the profound role of this pathway in muscle regenerative function in aging.

There is a paucity of information regarding the significance of ciliation in age-related diseases. Here, we demonstrate that an absence of ciliation is a cell-intrinsic change in aged MuSCs that diminishes their function. We find that loss of cilia contributes to a reduction in muscle regenerative capacity with aging. We note that the age-dependent loss in ciliation is not complete, but instead resembles the effect on MuSC ciliation of heterozygous deletion of IFT88 in young adult mice. This suggests that the MuSC primary cilium is highly sensitive to changes in levels of ciliary protein expression and thus particularly susceptible to aging-related changes. We show that the decrease in ciliation in aged MuSCs impairs Hedgehog (HH) signaling, suggesting an important role for HH in muscle regeneration. HH has previously been shown to play a role in myogenesis, especially during embryogenesis, as canonical HH signaling to somites is critical to the induction of myogenic factors such as MYOD1 and MYF5^29^, and the subsequent induction of slow twitch muscle fates^30^. In muscle progenitor cells, HH signaling has also been characterized as a pro-survival and proliferation factor^31^. Although there is limited *in vivo* evidence for the role of HH signaling in muscle regeneration, previous studies have shown that SMO inhibition by cyclopamine treatment of injured muscles results in muscle fibrosis and increased inflammation^32^. Additionally, in the context of aging, transduction of a *Shh*-expressing vector was shown to boost muscle repair to levels comparable to those found in much younger mice^33^. Here, we provide a previously unrecognized link of HH signaling to ciliation in MuSCs and show that HH signaling is important for the regenerative capacity of muscle *in vivo*. Furthermore, treatment with a small molecule that restores HH signaling in aged MuSCs improves endogenous stem cell function and significantly augments muscle regenerative capacity after injury of aged muscles.

Mesenchymal stem and progenitor cells are broadly ciliated and play an important role in maintaining stem cell fate by regulating lineage specification, proliferation and differentiation^34, 35, 36^. For example, in adipose tissue, preadipocytes are ciliated and their ablation leads to severe defects in white adipose tissue expansion^23^. Cilia also play a critical role in cartilage and bone development^37^. Finally, in skeletal muscle, although the function of cilia on MuSCs was not previously discerned, the presence of cilia was shown in another progenitor population, the fibroadipogenic progenitors^38, 39^, to play an important role in regulating its adipogenic fate in the context of injury and in a model of Duchenne Muscular Dystrophy^40^. Thus, ciliation of multiple progenitor cell populations is critical to tissue homeostasis and function.

Many signaling pathways are known to signal through the primary cilium. Here we implicate at least 5 ciliary signaling pathways in the regulation of aged MuSC proliferation, identified in a screen of a curated list of approximately 200 GPCR-targeting compounds. Additional ciliary signaling pathways not tested in this context may also play a role in muscle regeneration. This includes pathways activated in response to inflammation, since inflammatory signaling plays a critical role in almost all stages of muscle regeneration^13, 41, 42^ and is known to be transduced by the primary cilium in other cell contexts^43, 44^. Consistent with the potential role of the MuSC primary cilium in a multitude of signaling pathways, we show here that loss of ciliation in MuSCs results in a severe dysregulation of calcium and G coupled protein signaling. Indeed, how the primary cilium senses and integrates multiple ciliary signaling pathways into ciliary effector proteins to coordinate initiation of myogenesis during development remains an intriguing question^9^. Here, we demonstrate the profound functional importance of the primary cilium specifically in Hedgehog signaling in MuSC during regeneration of damaged tissue. Our findings provide fresh insights into the etiology of signaling dysfunction in aging and a potential therapeutic strategy to promote aged muscle regeneration.

## Acknowledgments

We apologize to those investigators whose important work we were unable to cite or describe owing to space constraints. We thank Barry Shearer and Patrick Eidam for their support in identifying the screening library. We thank FACS Core Facility in Stanford Lokey Stem Cell Research Building, and Stanford Veterinary Service Center (VSC) for technical support.

## Funding

This study was supported by the Baxter Foundation, the Li Ka Shing Foundation, and US National Institutes of Health (NIH) grants AG020961 (H.M.B.) and 5R01GM114276, 5U01CA199216, 5UL1TR00108502 (P.K.J). A.R.P is funded by a GSK Sir James Black Program for Drug Discovery Postdoctoral Fellowship. K.I.H is a Layton Family Fellow of the Damon Runyon Cancer Research Foundation (DRG-2210-14).

## Author contributions

A.R.P., K.I.H., J.P.K., A.C.H., P.K.J. and H.M.B. designed the experiments and wrote the manuscript. A.R.P., K.I.H., C.A.H., A.V.Y., N.Y., J.D., P.K. and B.C.G. performed the experiments and analyzed data. A.R.P. and K.I.H. contributed equally to the study. J.P.K., A.C.H. provided reagents necessary for the study. All authors discussed the results and commented on the manuscript.

## Data deposition

The data reported in this paper have been deposited in the Gene Expression Omnibus (GEO) database, https://www.ncbi.nlm.nih.gov/geo (accession no. GSE145297; GSE145312).

## Competing interests

J.P.K. and A.C.H. were employees of GSK during the time the work was performed.

## Data and materials availability

All data are available in the main text or the supplementary materials. Correspondence and requests for materials should be addressed to H.M.B.

