## Supplementary Material for "Ciliation of muscle stem cells is critical to maintain regenerative capacity and is lost during aging"

**This PDF file includes:**

Materials and Methods
Supplementary Figures S1-S5

### Materials and Methods

#### *Mice*

All studies were conducted in accordance with the GSK Policy on the Care, Welfare and Treatment of Laboratory Animals and were reviewed the Institutional Animal Care and Use Committee either at GSK or by the ethical review process at the institution where the work was performed. All experiments and protocols were performed in compliance with the institutional guidelines of Stanford University and Administrative Panel on Laboratory Animal Care (APLAC). Aged (>24 mo.) and young (2-4 mo.) C57BL/6 mice were obtained from Jackson Laboratory. Double-transgenic GFP/luc mice were generated as described previously [1]. Mouse transgenic strains were purchased from Jackson Laboratory (NOD-SCID No. 005557; $Pax7^{CreERT2}$ No. 017763; $IFT88^{flox/flox}$ *No.* 022409; *Rosa26-LSL-Luc* No. 005125, and $IFT88^{flox/flox}$ No. 022409). Double-transgenic $Pax7^{CreERT2}$*;Rosa26-LSL-Luc* were generated as described previously[2]. Double-transgenic $Pax7^{CreERT2}$;$IFT88^{flox/flox}$ ($IFT88^{f/f}$) were generated by crossing $Pax7^{CreERT2}$ mice obtained from Jackson Laboratory[3] and $IFT88^{f/f}$ obtained from Jackson Laboratory[4]. For $Pax7^{CreERT2}$*;Rosa26-LSL-Luc* and $Pax7^{CreERT2}$;$IFT88^{flox/flox}$ ($IFT88^{f/f}$) mice experiments, we treated 8-week-old male mice with five consecutive daily intraperitoneal injections of tamoxifen and performed intramuscular notexin injury one week after the last tamoxifen injection. We validated these genotypes by appropriate PCR-based strategies.

#### *Muscle stem cell isolation*

We isolated and enriched muscle stem cells as previously described [1, 2, 5, 6]. Briefly, hindlimb muscles were minced and digested using a collagenase and dispase solution by the MACs Dissociator (Miltenyi). Subsequently, single cells were depleted for hematopoietic lineage expressing and non-muscle cells ($CD45^-$/$CD11b^-$/$CD31^-$/$Sca1^-$) using a magnetic bead column (Miltenyi). The remaining $Lin^-$ cell mixture was then subjected to FACS analysis to sort for MuSCs co-expressing CD34 (Anti-Mouse CD34 eFluor 660, eBioscience, cat# 50-0341-82) and α7-integrin (Anti-a7 integrin antibody-APC conjugate, Ablabs) markers. We generated and analyzed flow cytometry scatter plots using FlowJo v10.0.

#### *Muscle stem cell transplantation*

We co-injected 250 GFP/luc IFT88$^{-/-}$ or control MuSCs into the *tibialis anterior* (TA) muscles of recipient NOD-SCID mice as previously described[1, 2, 5, 6]. GFP/luc IFT88$^{-/-}$ or control MuSCs were obtained by isolating MuSCs from IFT88$^{-/-}$ or control mice (2-4 mo.) post-tamoxifen injection and transducing with a luc-IRES-GFP lentivirus (GFP/luc virus) on day 1 of culture for a period of 24 hr before transplantation (see below "Muscle stem cell culture, treatment and lentiviral infection" section for details). We compared cells from different conditions by transplantation into the TA muscles of contralateral legs in the same mice. Four weeks after transplantation, mice were euthanized, and the TAs were collected for analysis.

*Bioluminescence imaging*

We performed bioluminescence imaging (BLI) using a Xenogen-100 system, as previously described [1, 2, 5, 6]. Briefly, we anesthetized mice using isofluorane inhalation and administered 120 μL D-luciferin (0.1 mmol $kg^{-1}$, reconstituted in PBS; Caliper LifeSciences) by intraperitoneal injection. We acquired BLI using a 60s exposure at F-stop=1.0 at 5 minutes after luciferin injection. Digital images were recorded and analyzed using Living Image software (Caliper LifeSciences). We analyzed images with a consistent region-of-interest (ROI) placed over each hindlimb to calculate a bioluminescence signal. We calculated a bioluminescence signal in radiance ($p\ s^{-1}\ cm^{-2}\ sr^{-1}$) value of $10^4$ to define an engraftment threshold. This radiance threshold of $10^4$ is approximately equivalent to the total flux threshold in p/s reported previously. This BLI threshold corresponds to the histological detection of one or more GFP+ myofibers [1, 2, 5, 6]. We performed BLI imaging every week after transplantation.

*Muscle injury*

We used an injury model entailing intramuscular injection of 10 or 20 μl of notexin (10 μg $ml^{-1}$; Latoxan) into the TA muscle or GA muscle respectively. When indicated, 3 days after injury SAG (175 μg/kg, Tocris, catalog # 4366) or vehicle control (PBS) was injected into the GA muscle as previously described [7]. We collected tissues at times indicated for analysis.

For *Pax7$^{CreERT2}$; Rosa26-LSL-Luc* mice experiments, we treated mice with five consecutive daily intraperitoneal injections of tamoxifen to activate luciferase expression under the control of the *Pax7* promoter. A week after the last tamoxifen injection, mice were subjected to intramuscular injection of 10 μl of notexin (10 μg $ml^{-1}$; Latoxan), which we designated as day 0 of the assay.

Three days later either SAG (175 µg/kg, Tocris, catalog # 4366) or vehicle control (PBS) was injected into the TA muscle. Bioluminescence was assayed at days 3, 7, 10 and 14 post-injury.

*Immunofluorescence staining and imaging*

We collected and prepared recipient TA muscle tissues for histology as previously described [2]. We fixed transverse sections from muscles using 4% PFA, blocked and permeabilized using PBS/1% BSA/0.1% Triton X-100 and incubated with anti-LAMININ (Millipore, clone A5, catalog # 05-206, 1:200) and then with AlexaFluor secondary Antibodies (Jackson ImmunoResearch Laboratories, 1:200) or wheat germ agglutinin-Alexa 647 conjugate (WGA, Thermo Fisher Scientific). We counterstained nuclei with DAPI (Invitrogen).

For myofibers and MuSCs, we performed fixation using 4% PFA, blocking and permeabilization using PBS/1% BSA/0.1% Triton X-100 and staining with primary antibodies anti-detyrosinated tubulin (abcam, catalog # ab48389, 1:100), anti-PAX7 (Santa Cruz Biotechnology, catalog # sc-81648, 1:50), anti-FOP (Abnova, catalog #H00011116-M01, 1:1000), anti-MyHC (MF20, Thermo Fisher Scientific, catalog # 14-6503-82, 1:500) and then with AlexaFluor secondary Antibodies (Jackson ImmunoResearch Laboratories, 1:500). We counterstained nuclei with DAPI (Invitrogen).

Confocal images of myofibers were acquired on a Marianas spinning disk confocal (SDC) microscopy (Intelligent Imaging Innovations) with a 40x/0.9 N.A. objective to capture multiple consecutive focal planes. Muscle transverse sections images were acquired on a KEYENCE BZ-X700 all-in-one fluorescence microscope (Keyence) with 20×/0.75 N.A. objectives. We analyzed the myofiber cross-sectional area using the Keyence Software that identified the fibers and segmented the fibers in the image to analyze the area of each fiber. For fiber area at least 10 fields of LAMININ-stained myofiber cross-sections encompassing over 400 myofibers were captured for each mouse. For percent ciliation, we analyzed young and aged MuSCs on at least 30 myofibers isolated from 5 independent mice using Intelligent Imaging Innovations software. Data analyses were blinded. The researchers performing the imaging acquisition and scoring were unaware of conditions given to sample groups analyzed.

*Hydrogel fabrication*

We used polyethylene glycol (PEG) hydrogels from PEG precursors, synthesized as described previously[6]. Briefly, we produced 12-kPa (Young's modulus) hydrogels in 1 mm thickness functionalized with laminin to cover the surface area of 12-well or 24-well culture plates.

*Myofiber isolation and culture*

EDL myofibers were isolated as previously described[8]. Briefly, the extensor digitum longus (EDL) was dissected and digested in 0.2% Type I Collagenase (Sigma) in DMEM at 37°C for 1 hour. Single myofibers were isolated by triturating the digested EDL muscle with polished Pasteur pipettes and then fixed.

*Cell culture*

Following FACs isolation, we resuspended MuSCs in myogenic cell culture medium containing DMEM/F10 (50:50), 15% FBS, 2.5 ng $ml^{-1}$ fibroblast growth factor-2 and 1% penicillin-streptomycin. We added 5 µM of SAG (Cayman Chemical Company, cat# 11914), 2 µM of GSA-10 (Sigma-Aldrich, cat# SML1171), 50 µM of Fluticasone (Sigma-Aldrich, cat# F9428), 5 µM of Cyclopamine (STEMCELL Technologies, cat#72072), or 0.1 µg/ml SHH (R&D Systems, cat# 461-SH-025) to MuSCs cultured on collagen coated dishes for the first 24h. The cells were then trypsinized and cells reseeded onto hydrogels for an additional 6 days of culture. All treatments were compared to their solvent (DMSO) vehicle control.

For $IFT88^{-/-}$ MuSCs transplant studies, we infected MuSCs with lentivirus encoding elongation factor-1α promoter–driven luc-IRES-GFP (GFP/luc virus) for 24h in culture as described previously[5]. Cells were assayed for GFP 48h post-infection using an inverted fluorescence microscope (Carl Zeiss Microimaging).

*Proliferation assays*

To assay proliferation, we seeded MuSCs on flat hydrogels at a density of 500 cells per $cm^2$ surface area. We counted cell number using a hemocytometer. We collected cells at indicated timepoints by incubation with 0.5% trypsin in PBS for 5 min at 37 °C and quantified them using a hemocytometer at least 3 times. Additionally, we used the VisionBlue Quick Cell Viability Fluorometric Assay Kit (BioVision, catalog # K303) as a readout for cell growth in culture. Briefly, we incubated MuSCs with 10% VisionBlue in culture medium for 4h, and measured fluorescence intensity on a fluorescence plate reader (Infinite M1000 PRO, Tecan) at Ex= 530–570nm,

Em=590-620nm. Data analyses were blinded, where researchers performing cell scoring were unaware of the treatment condition given to sample groups analyzed.

*Drug screen on aged MuSC proliferation*

To perform the drug screen, we seeded 200 MuSCs isolated from aged mice (>24 mo.) on collagen coated 96 well plates in myogenic cell culture medium containing DMEM/F10 (50:50), 15% FBS, 2.5 ng $ml^{-1}$ fibroblast growth factor-2 and 1% penicillin-streptomycin. We performed 5 replicates per condition plated on different replicate plates. Each plate contained vehicle, PGE2 (Cayman Chemical Company, cat# 14010, 10 ng/ml) and p38/MAPK inhibitor SB202190 (ApexBio, cat# A1632, 10 μM) controls. The chemical library against known GPCR receptors was derived from the Broad Institute Drug Repurposing Hub (https://clue.io/repurposing) and included both agonists and antagonists when possible [9]. Small molecules and peptides were added at a 2X concentration to achieve a final concentration of 10 μM. To avoid cell-washout effects, we did not change the medium throughout the assay. Proliferation was assayed at 7 days post-plating by using the VisionBlue Quick Cell Viability Fluorometric Assay Kit (BioVision, catalog # K303) as a readout for cell growth in culture. Briefly, we incubated live MuSCs with 10% VisionBlue in culture medium for 4h, and measured fluorescence intensity on a fluorescence plate reader (Infinite M1000 PRO, Tecan) at Ex= 530–570nm, Em=590-620nm. Additionally, at the endpoint cells were fixed using 4%PFA, stained with DAPI and imaged using the KEYENCE BZ-X700 all-in-one fluorescence microscope (Keyence) with a 4× objective. The entire plate was imaged and stitched using the Keyence Analysis Software, and cell confluency was determined for each well (**Fig. S1C**). Proliferation was normalized to the average value of vehicle treated for each plate. Data analyses were blinded, where researchers performing cell scoring were unaware of the treatment condition given to sample groups analyzed.

*Quantitative RT-PCR*

We isolated RNA from MuSCs using the RNeasy Micro Kit (Qiagen). We reverse-transcribed cDNA from total mRNA from each sample using the SensiFAST™ cDNA Synthesis Kit (Bioline). We subjected cDNA to RT-PCR using TaqMan Assays (Applied Biosystems) in an ABI 7900HT Real-Time PCR System (Applied Biosystems). We cycled samples at 95 °C for 10 min and then 40 cycles at 95 °C for 15 s and 60 °C for 1 min. To quantify relative transcript levels, we used

2−ΔΔCt to compare treated and untreated samples and expressed the results relative to *Gapdh (*Thermo Fisher Scientific, Cat # Mm01162710_m1).

TaqMan Assays (Applied Biosystems) were used to quantify *Gli2 (*Thermo Fisher Scientific, Cat # Mm01293117_m1 and *Smo* (Thermo Fisher Scientific, Cat # Mm01162710_m1) in samples according to the manufacturer instructions with the TaqMan Universal PCR Master Mix reagent kit (Applied Biosystems). Transcript levels were expressed relative to *Gapdh levels.* For Taqman qPCR, multiplex qPCR enabled target signals (FAM) to be normalized individually by their internal *Gapdh* signals (VIC).

*Flow cytometry*

We assayed EdU as a readout of proliferation for MuSCs after 7 days in culture on hydrogels, after an initial acute (24 hr) treatment of vehicle (DMSO), SAG or cyclopamine, or of control and $IFT88^{-/-}$ MuSCs. EdU incorporation was assessed using the Click-iT™ EdU Pacific Blue™ Flow Cytometry Assay Kit (Thermo, C10418) according to manufacturer's recommendation. Briefly, we incubated MuSCs for 1h with EdU for a final concentration of 20 µM after 7 days of proliferation on hydrogels. Cells were trypsinized and fixed in 4% PFA/PBS for 15 min at room temperature, followed by 2 rinses in 1%BSA/PBS. Cells were permeabilized in cold methanol for 10 min. Cells were kept in 1%BSA/PBS for 30min at room temperature, followed by 2 rinses in 1%BSA/PBS. Samples were then incubated in freshly made Click-iT reaction cocktail for 30min at room temperature, followed by 1 rinse in 3%BSA/PBS and 1 rinse in PBS. Samples were then incubated in 7AAD in PBS for 30min at room temperature followed by 1 rinse in PBS. We analyzed the cells on a FACS LSR II cytometer using FACSDiva software (BD Biosciences).

*In vivo muscle force measurement*

The peak isometric torque (N•mm) of the ankle plantarflexors was assessed as previously described[10, 11]. Briefly, the foot of anesthetized mice was placed on a footplate attached to a servomotor (model 300C-LR; Aurora Scientific). Two Pt-Ir electrode needles (Aurora Scientific) were inserted percutaneously over the tibial nerve, just posterior/posterior-medial to the knee. The ankle joint was secured at a 90° angle. The peak isometric torque was achieved by varying the current delivered to the tibial nerve at a frequency of 200 Hz and a 0.1-ms square wave pulse. We performed three tetanic measurements on each muscle, with 1 min recovery between each measurement. Data were collected with the Aurora Scientific Dynamic Muscle Data Acquisition and Analysis Software.

*RNA-Seq*

For RNA-seq, α7-integrin$^+$CD34$^+$ MuSCs were isolated as described above. RNA was isolated using Qiagen RNAEasy Micro kit from 5,000–10,000 cells and cDNA generated and amplified using NuGEN Ovation RNA-Seq System v2 kit. Libraries were constructed from cDNA with the TruSEQ RNA Library Preparation Kit v2 (Illumina) and sequenced to 30-40×$10^6$ × 75-bp reads per sample on a NextSeq 550 from the Stanford Functional Genomics Facility.

*RNA-Seq Analysis*

For the RNA-Seq analysis, bcbio-nextgen framework (https://bcbio-nextgen.readthedocs.io/) was used (version: 1.1.8-b): RNA sequences were aligned against the Mus musculus genome (mm10) using STAR[12]. RSEM[13] or Salmon[14] was used for calling transcripts and calculating transcripts per million (TPM) values as well as total counts. A counts matrix containing the number of counts for each gene and each sample was obtained. This matrix was analyzed by DESeq to calculate statistical analysis of significance[15] of genes between samples. The data reported in this paper have been deposited in the Gene Expression Omnibus (GEO) database, https://www.ncbi.nlm.nih.gov/geo (accession no. GSE145297; GSE145312).

*Statistical analyses*

We performed cell culture experiments in at least three independent experiments where three biological replicates were pooled in each. In general, we performed MuSC transplant experiments in at least two independent experiments, with at least 3-5 total transplants per condition. We used a paired t-test for experiments where control samples were from the same experiment *in vitro* or from contralateral limb muscles *in vivo*. A non-parametric Mann-Whitney test was used to determine the significance difference between untreated vs treated groups using $\alpha$=0.05. ANOVA or multiple t-test was performed for multiple comparisons with significance level determined using

Bonferroni correction or with Fisher's test as indicated in the figure legends. Unless otherwise described, data are shown as the mean ± s.e.m.

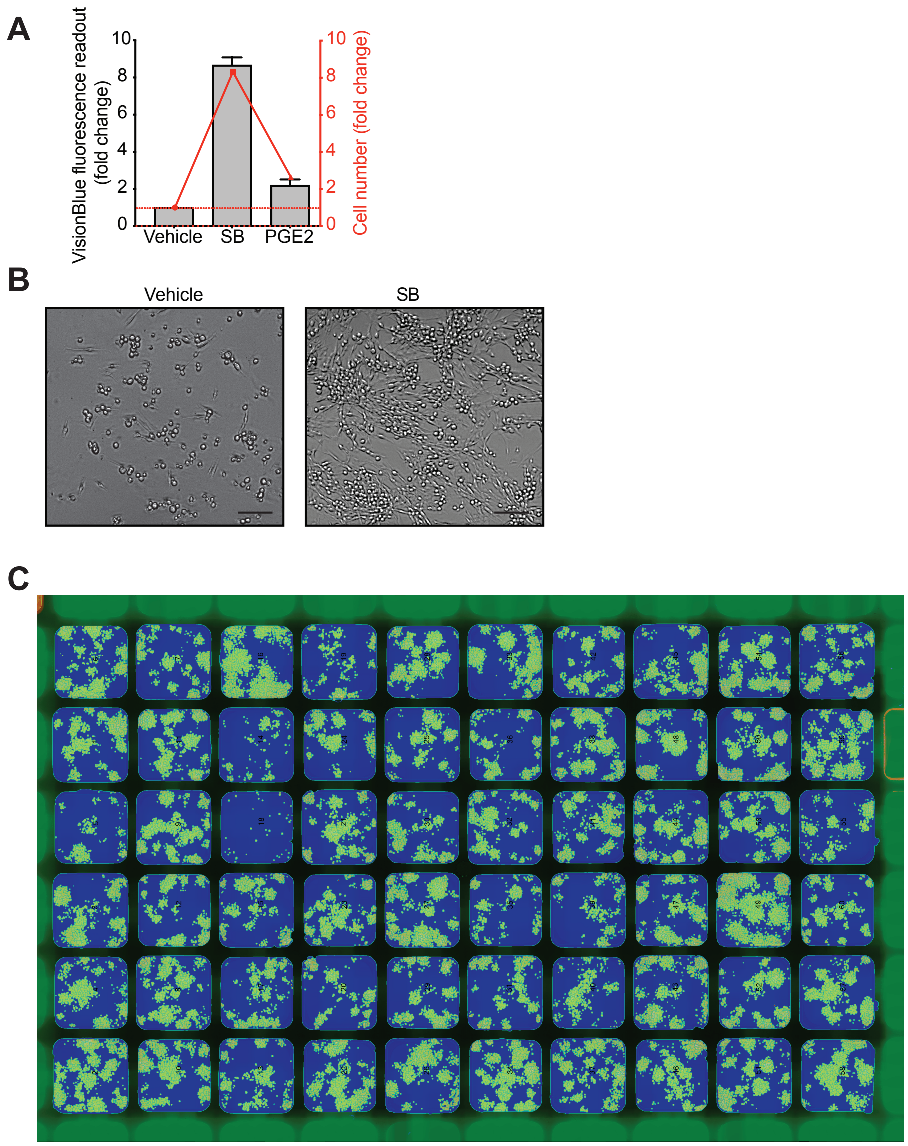


**Figure S1. Drug screen in aged MuSCs to assay proliferation**

**(A)** Comparison of VisionBlue fluorescence readout and cell number. MuSCs (α7+CD34+) were isolated and plated on collaged coated 96 well plates. VisionBlue was added to all wells and fluorescence was measured 4 hours later using a Tecan plate reader. Cells were then fixed and stained with DAPI. The entire well area was imaged, and cell number was calculated based on DAPI staining. Cell number and VisionBlue fluorescence readout were normalized to vehicle treated. Treatments using SB202190 (SB) and Prostaglandin E2 (PGE2) were used as positive controls. **(B)** Representative image of MuSCs treated with vehicle or SB. Bar=50 µm **(C)** Representative image of a 96-well plate with different drug treatments. Means+s.e.m.

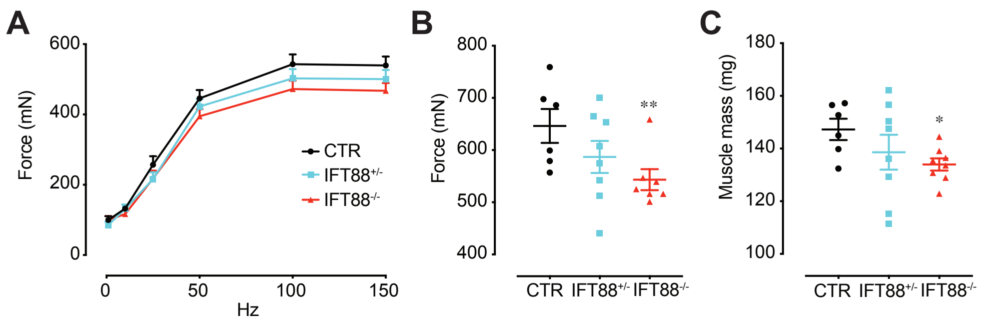


**Figure S2. Loss of cilia in MuSCs impairs muscle regeneration and strength recovery**

**(A)** Force-frequency curves of CTR, IFT88$^{+/-}$ and IFT88$^{-/-}$ mice 2 weeks post-injury. **(B)** Plantar flexion tetanic isometric force of CTR, IFT88$^{+/-}$ and IFT88$^{-/-}$ mice on day 14 post-injury (absolute levels). **(C)** Gastrocnemius muscle mass 2 weeks post-injury. *P<0.05, **P<0.01. Mann Whitney test performed for each condition compared to control (**B,C**). Means+s.e.m.

A

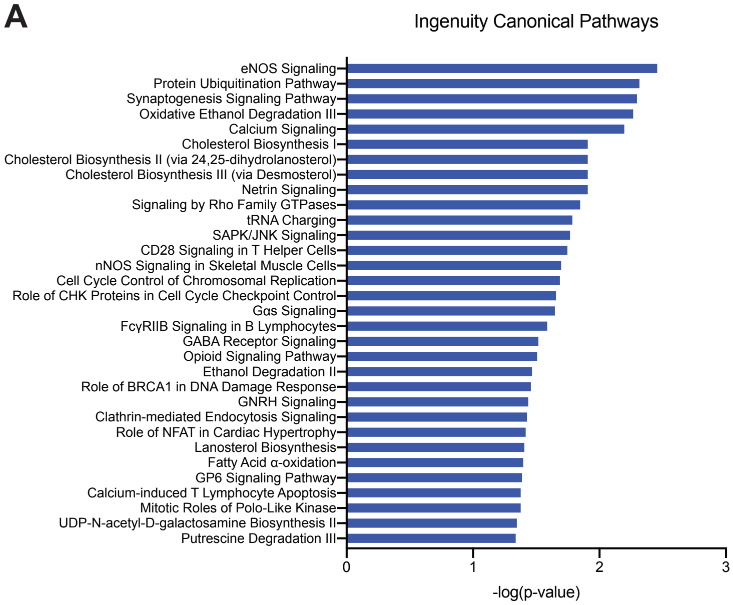


**Figure S3. Transcriptome analysis of control and IFT88$^{-/-}$ MuSCs**

**(A)** Enriched canonical pathways of the differentially expressed genes in IFT88$^{-/-}$ MuSCs indicated by Ingenuity Pathway Analysis (IPA).

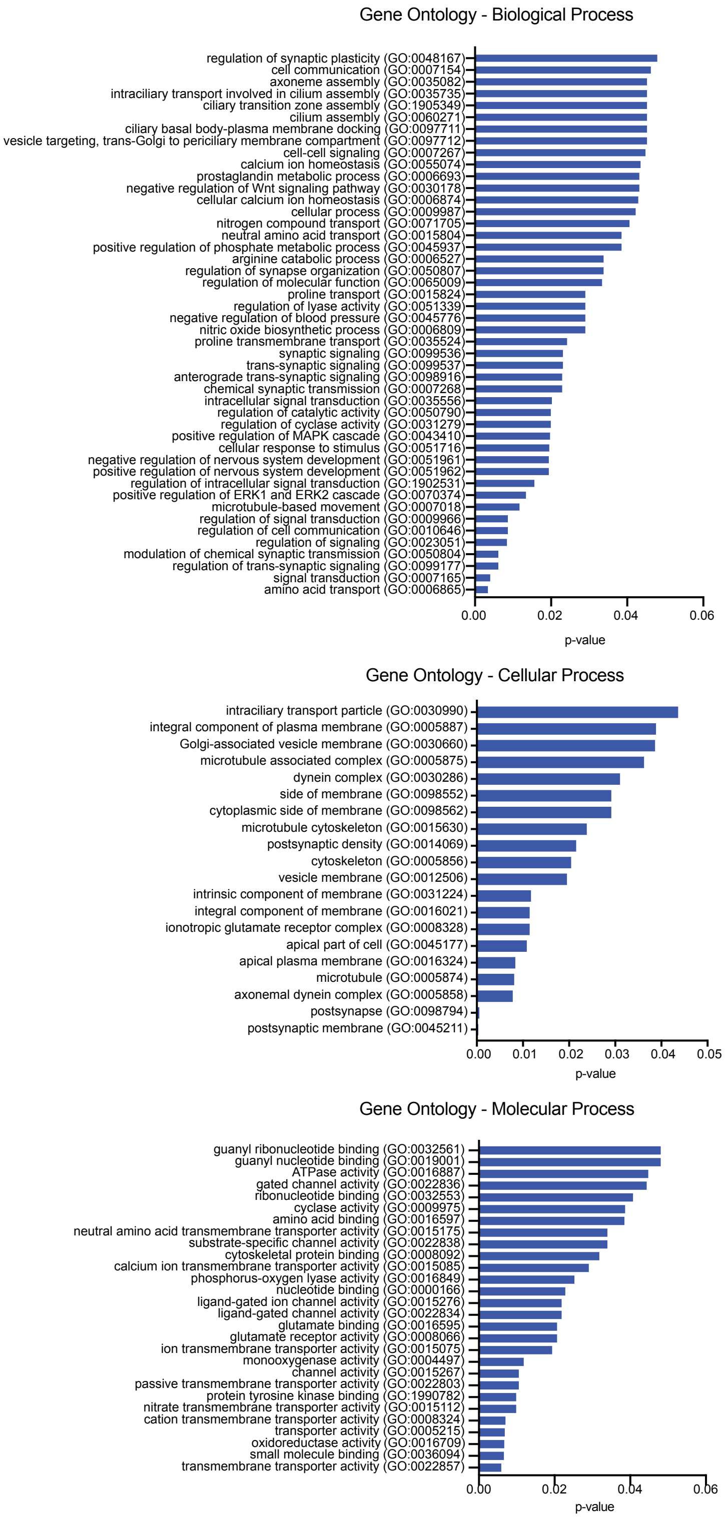
Gene Ontology - Biological Process
regulation of synaptic plasticity (GO:0048167)
cell communication (GO:0007154)
axoneme assembly (GO:0035082)
intraciliary transport involved in cilium assembly (GO:0035735)
ciliary transition zone assembly (GO:1905349)
cilium assembly (GO:0060271)
ciliary basal body-plasma membrane docking (GO:0097711)
vesicle targeting, trans-Golgi to periciliary membrane compartment (GO:0097712)
cell-cell signaling (GO:0007267)
calcium ion homeostasis (GO:0055074)
prostaglandin metabolic process (GO:0006693)
negative regulation of Wnt signaling pathway (GO:0030178)
cellular calcium ion homeostasis (GO:0006874)
cellular process (GO:0009987)
nitrogen compound transport (GO:0071705)
neutral amino acid transport (GO:0015804)
positive regulation of phosphate metabolic process (GO:0045937)
arginine catabolic process (GO:0006527)
regulation of synapse organization (GO:0050807)
regulation of molecular function (GO:0065009)
proline transport (GO:0015824)
regulation of lyase activity (GO:0051339)
negative regulation of blood pressure (GO:0045776)
nitric oxide biosynthetic process (GO:0006809)
proline transmembrane transport (GO:0035524)
synaptic signaling (GO:0099536)
trans-synaptic signaling (GO:0099537)
anterograde trans-synaptic signaling (GO:0098916)
chemical synaptic transmission (GO:0007268)
intracellular signal transduction (GO:0035556)
regulation of catalytic activity (GO:0050790)
regulation of cyclase activity (GO:0031279)
positive regulation of MAPK cascade (GO:0043410)
cellular response to stimulus (GO:0051716)
negative regulation of nervous system development (GO:0051961)
positive regulation of nervous system development (GO:0051962)
regulation of intracellular signal transduction (GO:1902531)
positive regulation of ERK1 and ERK2 cascade (GO:0070374)
microtubule-based movement (GO:0007018)
regulation of signal transduction (GO:0009966)
regulation of cell communication (GO:0010646)
regulation of signaling (GO:0023051)
modulation of chemical synaptic transmission (GO:0050804)
regulation of trans-synaptic signaling (GO:0099177)
signal transduction (GO:0007165)
amino acid transport (GO:0006865)
0.00
0.02
0.04
0.06
p-value
Gene Ontology - Cellular Process
intraciliary transport particle (GO:0030990)
integral component of plasma membrane (GO:0005887)
Golgi-associated vesicle membrane (GO:0030660)
microtubule associated complex (GO:0005875)
dynein complex (GO:0030286)
side of membrane (GO:0098552)
cytoplasmic side of membrane (GO:0098562)
microtubule cytoskeleton (GO:0015630)
postsynaptic density (GO:0014069)
cytoskeleton (GO:0005856)
vesicle membrane (GO:0012506)
intrinsic component of membrane (GO:0031224)
integral component of membrane (GO:0016021)
ionotropic glutamate receptor complex (GO:0008328)
apical part of cell (GO:0045177)
apical plasma membrane (GO:0016324)
microtubule (GO:0005874)
axonemal dynein complex (GO:0005858)
postsynapse (GO:0098794)
postsynaptic membrane (GO:0045211)
0.00
0.01
0.02
0.03
0.04
0.05
p-value
Gene Ontology - Molecular Process
guanyl ribonucleotide binding (GO:0032561)
guanyl nucleotide binding (GO:0019001)
ATPase activity (GO:0016887)
gated channel activity (GO:0022836)
ribonucleotide binding (GO:0032553)
cyclase activity (GO:0009975)
amino acid binding (GO:0016597)
neutral amino acid transmembrane transporter activity (GO:0015175)
substrate-specific channel activity (GO:0022838)
cytoskeletal protein binding (GO:0008092)
calcium ion transmembrane transporter activity (GO:0015085)
phosphorus-oxygen lyase activity (GO:0016849)
nucleotide binding (GO:0000166)
ligand-gated ion channel activity (GO:0015276)
ligand-gated channel activity (GO:0022834)
glutamate binding (GO:0016595)
glutamate receptor activity (GO:0008066)
ion transmembrane transporter activity (GO:0015075)
monooxygenase activity (GO:0004497)
channel activity (GO:0015267)
passive transmembrane transporter activity (GO:0022803)
protein tyrosine kinase binding (GO:1990782)
nitrate transmembrane transporter activity (GO:0015112)
cation transmembrane transporter activity (GO:0008324)
transporter activity (GO:0005215)
oxidoreductase activity (GO:0016709)
small molecule binding (GO:0036094)
transmembrane transporter activity (GO:0022857)
0.00
0.02
0.04
0.06
p-value

**Figure S4**

**Figure S4. Transcriptome analysis of control and IFT88$^{-/-}$ MuSCs**

Enriched biological, cellular and molecular processes of the differentially expressed up-regulated genes in the SAG-treated aged MuSCs indicated by Gene Ontology.

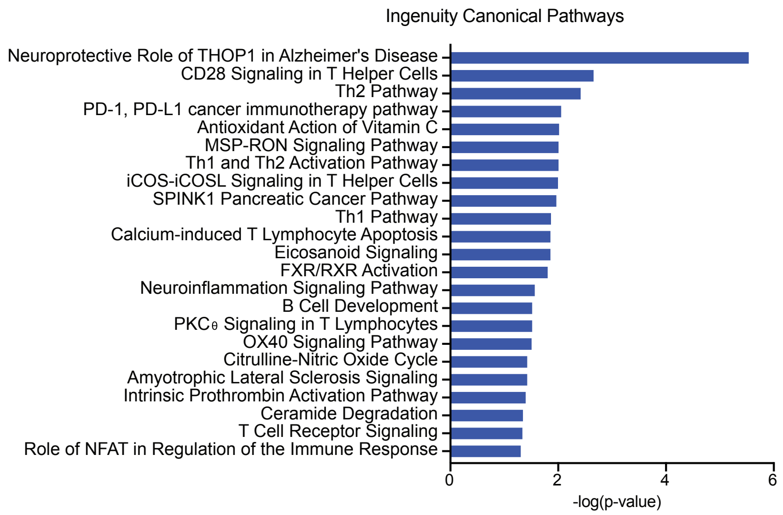


**Figure S5. Transcriptome analysis of SAG-treated aged MuSCs**

**(A)** Enriched canonical pathways of the differentially expressed genes in the SAG-treated aged MuSCs indicated by Ingenuity Pathway Analysis (IPA).
